# The *Magnaporthe oryzae* Effector AvrPii Attenuates Rice Immunity by Targeting the Calcium-Associated ANK Protein AVIN8

**DOI:** 10.64898/2026.05.13.725024

**Authors:** Wei Yao, Fuhui Yang, Xiaolong Zhou, Qianru Chen, Jinjun Lin, Yuying Zou, Chenyang Xu, Bei He, Dandan Zhu, Sisi Ji, Xionglun Liu, Jinling Liu

**Author notes:** Corresponding author: Jinling Liu,; Xionglun Liu.

## Abstract

Rice blast, caused by the fungus *Magnaporthe oryzae*, poses a severe threat to global rice production. Although numerous *M. oryzae* effectors have been identified, the molecular mechanisms remain poorly understood. The effector AvrPii triggers ETI upon recognition by NLR receptor Pii and has been shown to target host exocytosis and oxidative metabolic pathways to suppress immunity. Here, we identify AVIN8 (<u>Av</u>rPii-<u>in</u>teracting protein 8) as a new target of AvrPii. AVIN8 is an ankyrin-repeat (ANK) protein localized to the plasma membrane and nuclear envelope, with sequence similarity to known ANK calcium channels. *AVIN8* expression is induced during early *M. oryzae* infection. Overexpression of *AVIN8* enhances blast resistance, whereas its silencing increases susceptibility. AVIN8 promotes chitin-triggered ROS production in a Ca^2+^-dependent manner. In contrast, AvrPii impairs Ca^2+^ influx and chitin-triggered ROS burst. Our findings reveal a virulence strategy in which *M. oryzae* effector AvrPii hijacks the Ca^2+^ influx-associated ANK protein AVIN8 to suppress early calcium and PTI signaling in rice.

## Introduction

Plants employ a two-tiered innate immune system against a wide range of pathogens. The first layer, pattern-triggered immunity (PTI), is initiated by cell-surface localized pattern-recognition receptors (PRRs) that detect conserved microbial molecules, termed pathogen-associated molecular patterns (PAMPs). Adapted pathogens secrete effector proteins to suppress PTI, which can be recognized by intracellular receptors, such as nucleotide-binding leucine-rich repeat (NLR) proteins, triggering a stronger second layer immunity termed effector-triggered immunity (ETI) (Ngou et al., 2022; Jones et al., 2024). PTI and ETI are interconnected and share common downstream signaling, including calcium ions (Ca^2+^) influx, reactive oxygen species (ROS) burst, mitogen-activated protein kinase (MAPK) or calcium-dependent protein kinases (CDPK) signaling activation, and phytohormone signaling (Ngou et al., 2022). Among these events, Ca^2+^ influx and ROS burst are the earliest messengers and interconnected nodes in immunity activation (Köster et al., 2022). Ca^2+^ influx, mediated by diverse channels, such as CNGCs, GLRs, OSCAs, ANNs, NLRs, and ANK proteins, is critical for subsequent defense activation (Xu et al., 2022; Wang et al., 2024). Concurrently, ROS are produced by membrane-localized NADPH oxidases (RBOHs), and function as both signaling molecules and antimicrobial agents (Waszczak et al., 2018). Despite the critical role of Ca^2+^ influx in pathogen-induced ROS burst, the molecular interplay between these two signaling events remains elusive.

To counteract host immunity, pathogens have evolved sophisticated effector proteins to target the components of these immune signaling events, thereby suppressing host immunity and promoting infection (Hu et al., 2022; Kitazawa et al., 2022; Wang et al., 2022; Chen et al., 2025; Sun et al., 2025; Zhang et al., 2025; Zheng et al., 2025). Among the diverse virulence strategies, the blast fungus *Magnaporthe oryzae*, one of the most devastating fungal diseases of rice, employs two major types of effectors-apoplastic and cytoplasmic, to reprogram host immunity and metabolism (Oliveira-Garcia et al., 2024). Apoplastic effectors Slp1 and MoPMO9A sequester fungal chitin to prevent its recognition by the host receptor OsCEBiP, thereby blocking PTI (Mentalck et al., 2012; Martinez-D’Alto et al., 2023). Cytoplasmic effectors target diverse intracellular immune components, including the proteasome (Park et al., 2012), the exocyst complex (Fujisake et al., 2015; De la Concepcion et al., 2022), ROS metabolism (Singh et al., 2016; Han et al., 2021; Ning et al., 2022), ion channels (Shi et al., 2018), transcriptional regulators (Kim et al., 2020), hormone signaling (Liu et al., 2022; Liu et al., 2025), heavy metal-associated domain containing (HMA) proteins (Oikawa et al., 2020), and protein kinase (Wang et al., 2026). Notably, a subset of cytoplasmic effectors, termed avirulence (Avr) proteins, play dual roles contingent upon host genotype. They trigger ETI when recognized by specific intracellular resistance (R) receptors (typically NLRs), but otherwise act as virulence factors to suppress host defense. Although 13 *M. oryzae* Avr effectors have been molecularly identified (Oliveira-Garcia et al. 2024; Huang et al., 2025), the molecular mechanisms of most remain poorly understood.

Ankyrin-repeat (ANK) proteins are evolutionarily conserved (Wu et al., 2025) and play critical roles in immunity across multiple plant species, including Arabidopsis (Lu et al., 2003; Yang et al., 2012), rice (Chern et al., 2005; Wang et al., 2006; Zhang et al., 2010; Jiang et al., 2013; Mou et al., 2013) and wheat (Kolodziej et al., 2021). However, their precise biochemical and physiological functions in plant immune regulation remain largely elusive. Notably, recent studies revealed that some plant ANK proteins, such as ACD6 and Lr14a, function as calcium-permeable channels, unveiling a novel mechanism for ANK-mediated immune regulation (Chen et al., 2023; Yue et al., 2025). Despite the growing recognition of ANK proteins in immunity, their role as direct targets of pathogen effectors remains underexplored.

Among the characterized *M. oryzae* effectors, AvrPii is recognized by the NLR receptor Pii in rice, triggering ETI (Vo et al., 2019). Several host targets of AvrPii have been reported including exocyst subunit OsExo70-F2/F3 (Fujisake et al., 2015; De la Concepcion et al., 2022) and metabolic enzyme OsNADP-ME2 (Singh et al., 2016), yet its full target repertoire remains undefined. In this study, we identify a novel host target of AvrPii, designated AVIN8 (AvrPii-interacting protein 8). We demonstrate that AVIN8 is an ANK protein localized to the plasma membrane and nuclear envelope, and its ANK domain shares conserved residues with known calcium-permeable channels. *AVIN8* expression is induced during early infection, and functional analyses reveal that AVIN8 positively regulates blast resistance and enhances chitin-triggered immune response. Furthermore, AVIN8 promotes ROS generation in a Ca^2+^-dependent manner, whereas AvrPii suppresses this response. Thus, our findings reveal a virulence strategy in which the *M. oryzae* effector AvrPii hijacks the Ca^2+^ influx-associated ANK protein AVIN8 to dampen early calcium influx and immunity, thereby facilitating infection in rice.

## Materials and methods

### Plant materials and growth conditions

The rice (*Oryza sativa* L. *spp*. japonica) cultivars Nipponbare (NPB) and the *R* gene *Pii* near-isogenic line Pii-NIL were used in this study. Rice plants were grown in controlled greenhouse conditions at 28 °C with 70% relative humidity and a 12 h light/12 h dark photoperiod. For field experiments, plants were cultivated in Changsha, China, during typical growing season (March to July) under standard field management practices.

### Constructs preparation and rice genetic transformation

For overexpression construct, the full-length *AVIN8* cDNA was amplified and inserted into the *Xcm*I site of the binary expression vector pCXUN-HA (driven by the ubiquitin promoter) using a TA cloning strategy as previously described (Chen et al., 2009). For the RNAi construct, a 210-bp fragment derived from the 3′ untranslated region (3′ UTR) of *AVIN8* was selected to generate double-stranded RNA (dsRNA). The fragment was then cloned into the binary vector pANDA via Gateway® technology, following an established protocol (Li et al., 2024). Transgenic rice plants were generated in the NPB background using *Agrobacterium tumefaciens*-mediated transformation method.

### RNA extraction and qRT-PCR analysis

Total RNA was isolated from rice tissues using the Eastep® Super Total RNA Extraction Kit (Promega, Cat. #LS1040). The extracted RNA was then treated with DNase I (Promega, Cat. #LS1040) to remove genomic DNA contamination. First-strand cDNA was synthesized from 1 μg of total RNA using a reverse transcription kit (Promega, Cat. #A5001). The cDNA was diluted 10-fold, and 1 μL of the dilution product was used as template in a 20 μL reaction system containing SYBR qPCR Master Mix (Vazyme, Cat. #Q312-02). Quantitative PCR (qPCR) was performed on a CG-05 Real-Time PCR Detection System (Hangzhou Jingle Scientific Instrument Co., Ltd, China) under the following cycling conditions: initial denaturation at 95 °C for 30 s, followed by 40 cycles of 95 °C for 10 s and 60 °C for 30 s. The relative expression levels of targeted genes were calculated using the 2^-ΔΔCt^ method, with *Ubiquitin* (*Ubq*) as the internal reference. Two independent biological replicates, each consisting three technical replicates, were analyzed. Data are presented as the mean ± standard deviation (SD) for one representative biological experiment.

### Yeast two-hybrid assay

Yeast two-hybrid analysis was performed using the ProQuest™ Two-Hybrid System (Invitrogen) following the manufacturer’s instructions. Briefly, the full-length *AvrPii* cDNA was cloned into *SalI* and *NotI* sites of the bait vector pDBleu to generate BD fusion construct. Similarly, the full-length *AVIN8* cDNA was inserted into the *SmaI* and *NotI* sites of the prey vector pPC86 to produce AD fusion construct. Both constructs, along with their respective empty vectors as controls, were co-transformed into the *Saccharomyces cerevisiae* strain MAV203 using the standard lithium acetate/ polyethylene glycol method. Transformants were plated onto synthetic defined (SD) selection medium supplemented with optimal concentrations of 3-amino-1,2,4-triazole (3-AT) and incubated at 28 °C in darkness for 3-4 days. Protein-protein interactions were assessed based on yeast colony growth phenotypes under selective conditions.

### Bimolecular fluorescence complementation (BiFC) assay

The coding sequences of *AvrPii* and *AVIN8* were cloned into the *Bam*HI/*Xho*I sites of the vector KanII-SPYCE (carrying the C-terminal half of YFP, cYFP) and the *Sal*I/*Xho*I sites of the vector hygII-SPYNE (R)173 (carrying the N-terminal half of YFP, nYFP), respectively (Waadt et al., 2008). All constructs and corresponding empty vectors (EVs) were individually transformed into *Agrobacterium tumefaciens* strain EHA105. Bacterial suspensions harboring the following plasmid combinations of *AvrPii-nYFP*/ *cYFP-AVIN8, AvrPii-nYFP*/*cYFP-EV, nYFP-EV*/*cYFP-AVIN8, nYFP-EV/cYFP-EV* were co-infiltrated into leaves of *N. benthamiana* plants. After incubation for 48h, infiltrated leaf samples were harvested and imaged under a confocal laser-scanning microscope (Leica, TCS SP8). YFP fluorescence signals were detected using excitation/emission wavelength of 515 nm/525 nm.

### Subcellular localization

The full-length *AVIN8* cDNA was cloned into the sites of *Bgl*II and *Sal*I sites of the binary expression vectors pGDR, respectively (Goodin et al., 2000). The constructs were then transformed into *Agrobacterium* strain EHA105. The *Agrobacterium* suspensions harboring the respective constructs were infiltrated into *N. benthamiana* leaves. After 48 h of infiltration, leaf samples were collected and examined for fluorescence under a confocal microscope (Leica, TCS SP8). GFP and RFP signals were detected at 488 nm excitation/emission wavelengths of 488 nm/500-550 nm and 515 nm/580-630 nm, respectively.

### Rice blast inoculation and resistance evaluation

*M. oryzae* isolates were cultured on oatmeal agar plates (20 g/L oatmeal [Pure oat flakes, Seamild],14 g/L agar [Sangon Biotech, Cat. #A505255-0250]) at 28 °C in darkness for 7 days. Plates were then transferred to continuous light for 15-20 days to induce conidia. Conidia were harvested by gently scraping the culture surface with sterile 0.02% (v/v) Tween-20 solution, and the spore suspension was adjusted to the desired concentration for subsequent inoculations.

For in vitro detached leaf inoculation, leaf segments (7 cm in length) were excised from the second youngest fully expanded leaf of rice seedlings at 3-4 leaf stage. Segments were placed in Petri dishes with filter paper moistened with 10 μg/mL 6-benzylaminopurine solution. Two puncture wounds were made along the central vein on the adaxial side of each leaf. A 5 μL droplet of spore suspension (2×10^6^ spores/mL in 0.02% Tween-20) was applied to each wound. Dishes were sealed and incubated at 26-28 °C and 80% RH under continuous darkness for 24 h, followed by a 12 h light/12 h dark cycle for 5-7 days. Lesion length and area were measured to quantify disease severity.

For whole-plant spray inoculation, 3-4 leaf stage seedlings were uniformly sprayed with a spore suspension (5×10^5^ conidia/mL in 0.02% Tween-20 solution). After inoculation, plants were kept in darkness at high humidity for 24 h, then transferred to a growth chamber set at 28 °C, 85% RH, with a 12 h light/12 h dark photoperiod. Disease symptoms were assessed at 7 days post-inoculation.

For field nursery inoculation, germinated seeds were sown in a field nursery. At the 1-2 leaf stage, seedlings were inoculated by evenly scattering *M. oryzae*-infected rice straw (collected from previous epidemic seasons and stored dry) across the nursery to facilitate natural infection. Disease development was evaluated after 2-3 weeks.

### Fungal biomass quantification by qPCR

Total genomic DNA was extracted from blast-infected leaf tissues using standard CTAB method. The extracted DNA was then used as a template for qPCR to amplify the rice *Ubiquitin* gene and the *M. oryzae-*specific repetitive element *MoPot2*. Reactions were carried out in a 20 μL volume containing SYBR qPCR Master Mix (Vazyme, Cat. #Q312-02) on a CG-05 Real-Time PCR Detection System. The thermal cycling conditions were as follows: initial denaturation at 95 °C for 30 s, followed by 40 cycles of 95 °C for 10 s and 60 °C for 30 s. Relative fungal biomass was calculated using 2^-ΔΔCt^ method, with the rice *Ubiquitin* gene as an internal control. Two independent biological replicates, each with three technical repetitions, were analyzed. Data from one representative biological experiment are presented as mean ± SD. Statistical significance was evaluated using one-way ANOVA method.

### Detection of reactive oxygen species (ROS) accumulation

ROS burst was measured using luminol-based chemiluminescence assay as previously described (Liu et al., 2023). Briefly, 3-mm leaf sheath segments were excised from three-leaf stage seedlings and pre-incubated in sterile distilled water overnight. Segments were then transferred to a reaction buffer containing 8 nM chitin (Qingdao BZ Oligo Biotech, Cat. # SCN05), 10 nM flg22, 10 mM CaCl_2_, or 10 μM LaCl_3_ (calcium channel inhibitor), 20 µM L-012 (Wako, Cat. #120-04891) and 10 µg/mL of horseradish peroxidase (Sangon Biotech, Cat. #A600691-0100). Chemiluminescence was monitored continuously using a LumiStation 1800 microplate reader (Shanghai Flash Spectrum Biological Technology Co., Ltd., Shanghai, China), and readings were recorded at 30 s interval over 50 time points. Five independent biological replicates were performed. Data are presented as the means ± SD.

### Gene accessions

The gene accessions used in this study are referenced in the Rice Genome Annotation Project version 7 as following: *AVIN8* (Os09g33810), *Ubquitin* (Os03g13170), *PBZ1* (Os12g36880), *OsCEBiP* (Os03g04110), *OsCERK1* (Os08g42580). The information of primer pairs for these genes used in this study are listed in Table S1.

## Results

### AvrPii interacts with rice host protein AVIN8 at the cell membrane

To elucidate the mechanisms underlying AvrPii-mediated immune modulation in rice, we performed a Y2H screening and identified a rice protein, designated AVIN8 (<u>Av</u>rPii-<u>in</u>teracting protein 8), as a candidate interactor. A subsequent Y2H assay using the full-length AVIN8 confirmed the specific interaction. Yeast co-transformed with BD-AvrPii and AD-AVIN8 grew on selective medium lacking leucine, tryptophan, and histidine (SD/-Leu-Trp-His) and supplemented with 20 mM 3-AT, whereas control combinations (BD-AvrPii/AD-EV, BD-EV/AD-AVIN8, and BD-EV/AD-EV) failed to grow (Fig.1A). To examine the interaction *in-planta*, we employed a BiFC assay. Co-expression of AvrPii-cYFP and nYFP-AVIN8 in *N. benthamiana* leaves resulted in strong YFP fluorescence at both the plasma membrane and nuclear envelope. No fluorescence was observed in the control combinations (nYFP-AVIN8/cYFP-EV, or nYFP-EV/AvrPii-cYFP) (Fig.1B). Therefore, these results demonstrated that AvrPii directly interacts with AVIN8 and this interaction occurs at the cell membrane.

**Figure 1.**
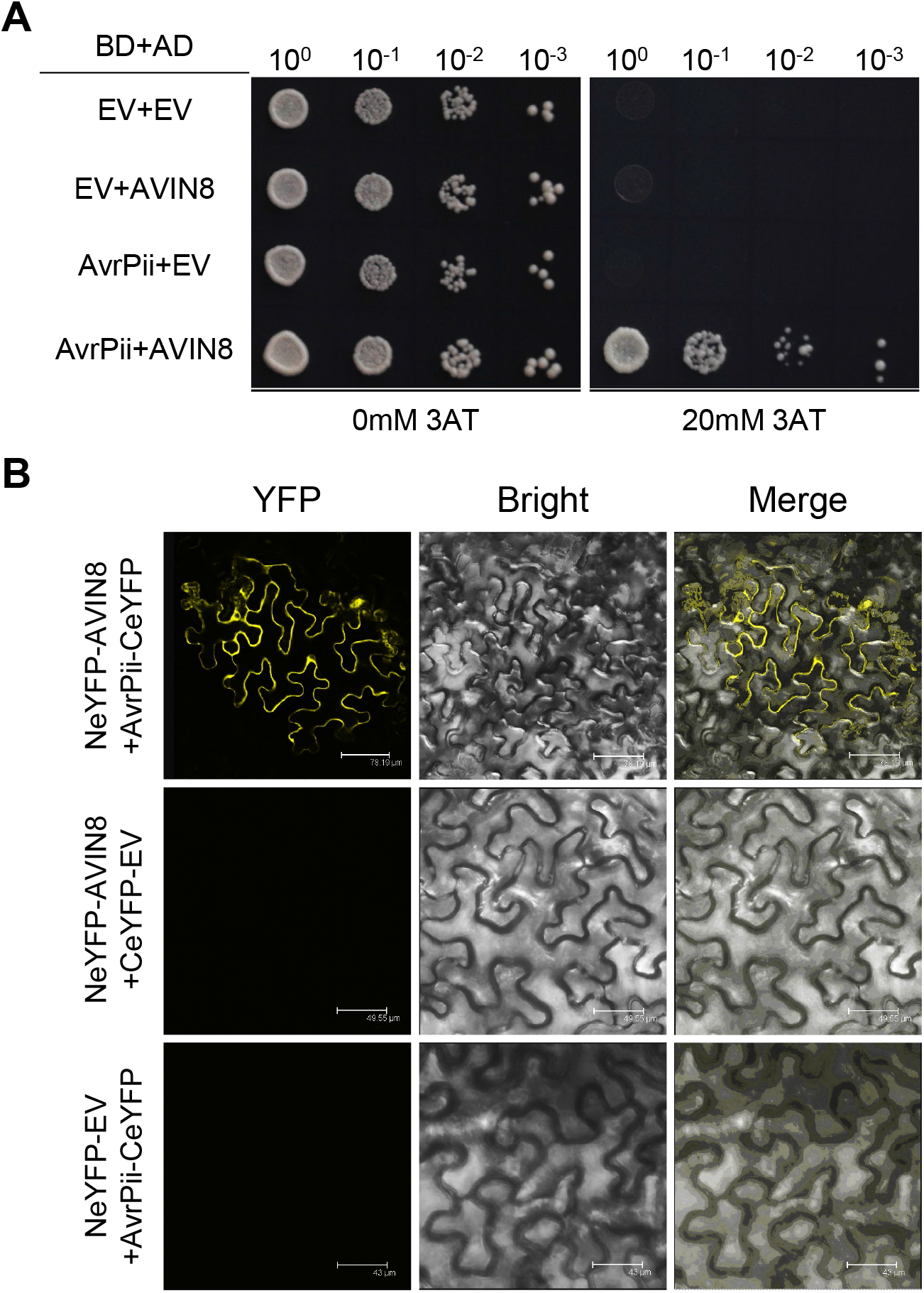
AvrPii interacts with AVIN8 in vitro and in planta. A. AvrPii interacts with AVIN8 in yeast. The top row indicates the dilution series; the left column shows the growth of yeast on SD-Leu-Trp-His+0 mM 3AT medium, and the right column shows the growth of yeast on SD-Leu-Trp-His+ 20 mM 3AT screening medium. EV: empty vector. B. The BiFC assay for the interaction between AvrPii and AVIN8 in-planta. AvrPii and AVIN8 were respectively fused with C-terminal eYFP (CeYFP) and N-terminal eYFP (NeYFP). Scale bars are shown in the figure.

### AVIN8 is a cell membrane localized ankyrin repeat protein

To characterize the function of AVIN8, we first analyzed its conserved domain using the NCBI Conserved Domain Database. A BLASTp search identified three conserved regions: an N-terminal GerD domain (residues 33-99)-a domain in GerD protein for nutrient-sensing involved in spore germination in *Bacillus* species (Li et al., 2008), a central STI1 domain (residues 135-178)-a conserved motif in the Sti1 proteins that are cochaperones associated with Hsp70/Hsp90 foldosome system (Schmid et al., 2012), and a C-terminal ankyrin-repeat (ANK) domain (residues 203-323). The presence of this canonical ANK domain confirms that AVIN8 is a member of the ankyrin-repeat protein family (Fig.2A).

**Figure 2.**
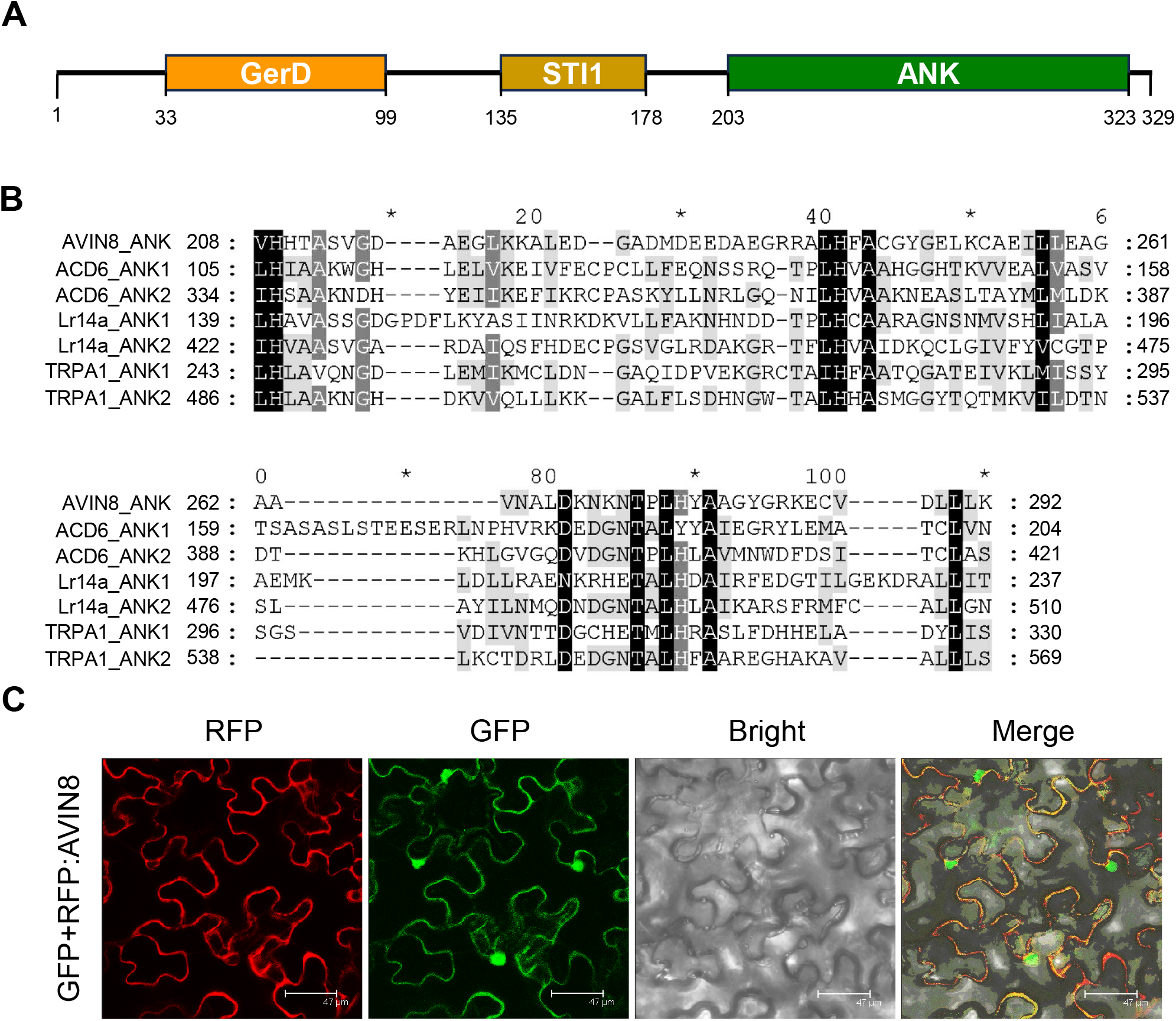
*AVIN8* encodes a cell membrane localized ankyrin-repeat protein. A. The protein encoding structure of AVIN8. GerD is a core domain in GerD protein from Bacillus species. STI1 is a conserved domain in the Sti1 proteins. ANK is a conserved domain composed of tandem repeats of a 30-34 amino-acid motif. B. Amino acid sequence alignments of ANK domains of AVIN8 and its homologs. C. Subcellular localization of AVIN8 in *N. benthamiana* leaf cells. GFP: green fluorescence; RFP: red fluorescence; GFP-AVIN8 is N-terminal GFP fused AVIN8 protein, RFP-AVIN8 is N-terminal RFP fused AVIN8 protein; Yellow fluorescence is merged color of GFP and RFP fluorescence. The bar represents 47 μm.

Amino acid sequence alignment of the ANK domains revealed that, AVIN8 shares 17.27-50.59% overall similarity with the known calcium-channel proteins ACD6, Lr14a and TPRA1, with the highest similarity (50.59%) to TRPA1_ANK2 domain (Fig. S1A). Key amino acids residues are fully conserved among them (Fig. 2B). Phylogenetic analysis further indicated that the ANK domain of AVIN8 clustered in one clade with TRPA1_ANK1, TRPA1_ANK2 and Lr14a_ANK1 (Fig. S1B), supporting a potential functional association between AVIN8 and calcium-channel regulators.

Furthermore, subcellular localization was examined by transiently expressing RFP-tagged AVIN8 in *N. benthamiana* leaves. RFP-AVIN8 fluorescence was observed at both the plasma membrane and nuclear envelope. When co-expressed with a free GFP control, which labels the entire cell, the fluorescence of RFP-AVIN8 only overlapped with GFP signal specifically at the plasma membrane and nuclear envelope, but not within the nucleus (Fig. 2C), confirming that AVIN8 is a dual-membrane localized protein.

### *AVIN8* expression is induced early during blast infection

RT-qPCR analysis revealed that *AVIN8* is constitutively expressed across all examined rice tissues, with the highest transcript levels detected in stems (Fig. 3A). To investigate its response to pathogen challenge, we monitored *AVIN8* transcription in an isogenic rice line with *R* gene *Pii* and NPB without Pii after *M. oryzae* inoculation. In *Pii* isogenic line, *AVIN8* expression showed a transiently low level of increase at 4h, followed by a decline by 48h in mock treatment. While upon *M. oryzae* infection, *AVIN8* exhibited a rapid induction with higher transcription level than those in mock controls from 4h to 72h post-inoculation (hpi) (Fig. 3B). Similarly, *AVIN8* displayed a consistent induced expression pattern in the background of NPB without *R* gene *Pii* (Fig. 3C). This induction was observed in both *Pii*-containing and *Pii*-lacking backgrounds, indicating it is independent to *R* gene recognition.

**Figure 3.**
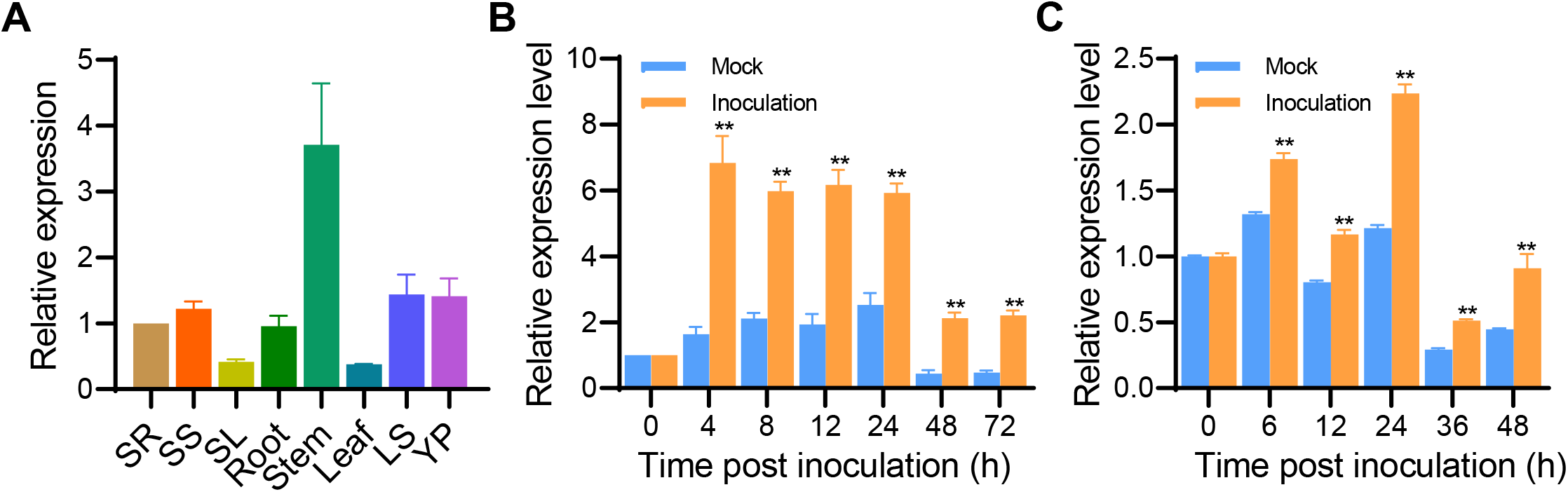
*AVIN8* is constitutively expressed and transcriptionally induced by *M. oryzae* infection. A. Transcription pattern of *AVIN8* gene in rice tissues. SR: seedling root; SS: Seedling sheath; SL: seedling leaf; LS: leaf sheath at the flowering time. YP: Young panicle before flowering. B. The expression dynamic of *AVIN8* after inoculated with blast strain 193-1-1 in the background of blast *R* gene *Pii* isogenic line. C. The expression dynamic of *AVIN8* after inoculated with blast strain 318-2 in background of NPB without blast R gene *Pii*. Data were analyzed by one-way ANOVA method with significance level marked with \**P* < 0.05; \*\**P* < 0.01.

### AVIN8 positively regulates rice blast resistance

To determine the biological function of AVIN8 in rice immunity, we generated *AVIN8* overexpression (OE) and RNA interference (RNAi) transgenic lines (Fig. S2). Disease resistance was first assessed using an *in vitro* punch inoculation method infected with *M. oryzae* strain 318-2. Compared with the wild-type NPB control, *AVIN8-*OE lines (OE-1 and OE-2) exhibited significantly enhanced resistance, showing shorter length and reduced relative fungal biomass (Fig. 4A-C). In contrast, *AVIN8*-RNAi lines displayed increased susceptibility, with longer lesions and higher fungal biomass relative to NPB (Fig. 4D-F). These phenotypes were further confirmed by whole-plant spray inoculation and field-nursery resistance evaluation under natural infection conditions. Overexpression of *AVIN8* significantly enhanced blast resistance, while silencing of *AVIN8* increased the susceptibility to blast fungal in both spray and field inoculation conditions (Fig. 4G-J). These results demonstrate that AVIN8 acts as a positive regulator of rice blast resistance.

**Figure 4.**
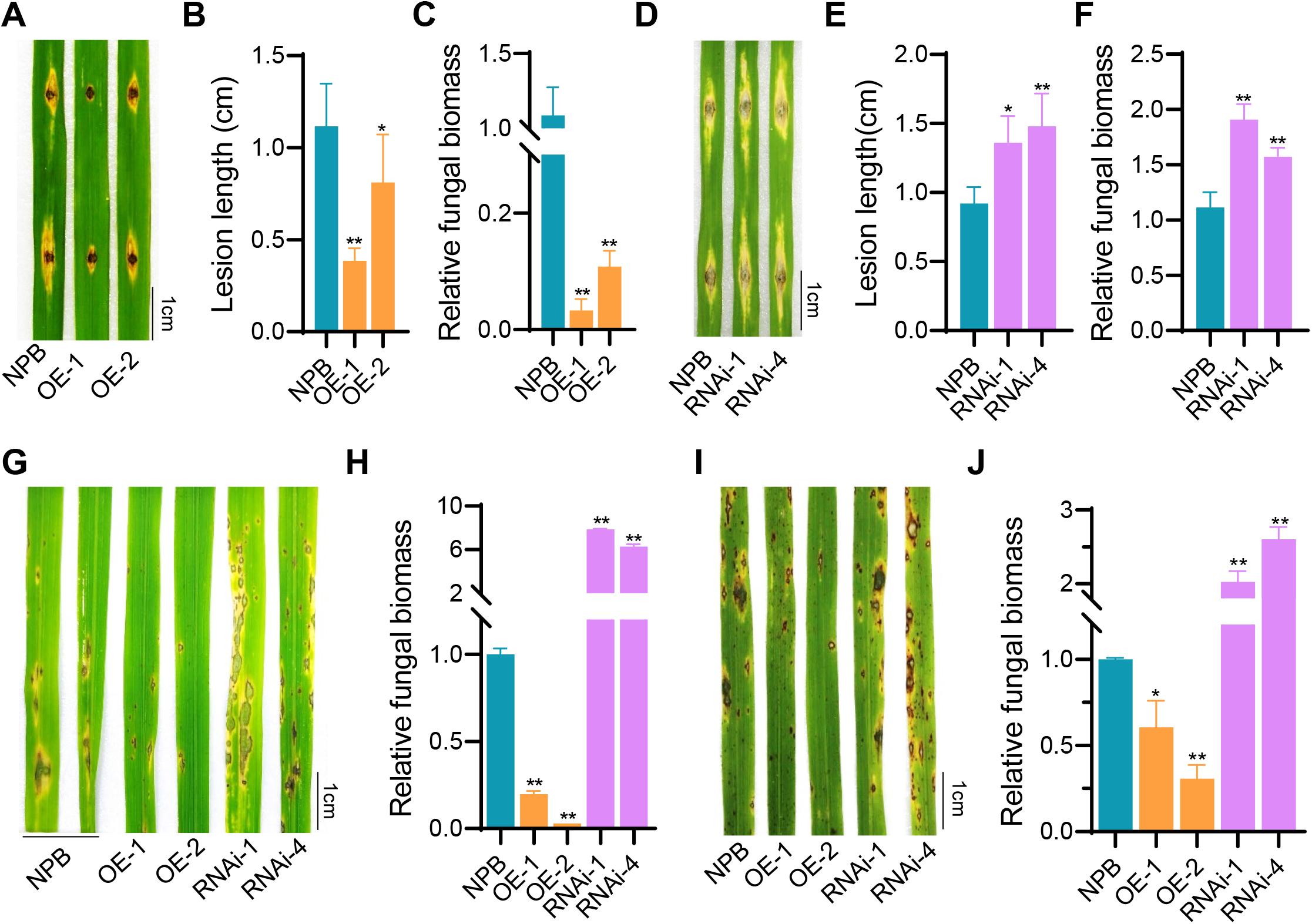
AVIN8 positively regulates disease resistance against *M. oryzae*. A-C. Disease phenotype (A), lesion length (B) and relative fungal biomass (C) of *AVIN8*-overexpressing (OE) transgenic rice inoculated by detached leaf punch method with *M. oryzae* race 318-2. OE-1 and OE-2 are two independent lines; NPB: wild-type control; 318-2: *M. oryzae* strain with AvrPii gene. The bar represents 1 cm in length. D-F. Disease phenotype (D), lesion length (E) and relative fungal biomass (F) of *AVIN8*-silencing transgenic rice inoculated using the whole-plant spray method with *M. oryzae* race 318-2. RNAi-1 and RNAi-4 represent two independent RNAi lines. The bar represents 1 cm in length. G-H. Disease phenotype (G) and relative fungal biomass (H) of *AVIN8* transgenic rice lines inoculated by spray method with *M. oryzae* race 318-2. I-J. Disease phenotype (I) and relative fungal biomass (J) of *AVIN8* transgenic rice lines in field nursery inoculation. The scale bar represents 1 cm in length. Significance level was analyzed by one-way ANOVA method (\**P* < 0.05; \*\**P* < 0.01).

### AVIN8 modulates early PTI signaling

To decipher the signaling pathway underlying AVIN8-mediated immunity, we examined the transcriptional dynamics of defense-related genes after blast inoculation. The results showed that the expression of pathogenesis-related (*PR*) genes *PBZ1* was significantly up-regulated in *AVIN8-*OE line but down-regulated in AVIN8-RNAi line during early infection (0-12hpi), with a peak at 6hpi (Fig.5A), suggesting that AVIN8 positively regulates early immune signaling against *M. oryzae*.

To further identify the early pathways influenced by AVIN8, we analyzed the expression of *OsCEBiP* and *OsCERK1*, two core receptors required for chitin perception and PTI activation (Gong et al., 2020). The results showed that both genes were rapidly induced in wild-type NPB and *AVIN8-*OE line during early *M. oryzae* infection. Overexpression of *AVIN8* significantly upregulated *OsCEBiP* expression compared with NPB, but did not affect *OsCERK1* expression. In contrast, silencing *AVIN8* markedly suppressed the induced expression of both *OsCEBiP* and *OsCERK1* (Fig. 5B, C). These results suggest that AVIN8 contributes to chitin-triggered PTI, likely through its association with chitin receptors.

**Figure 5.**
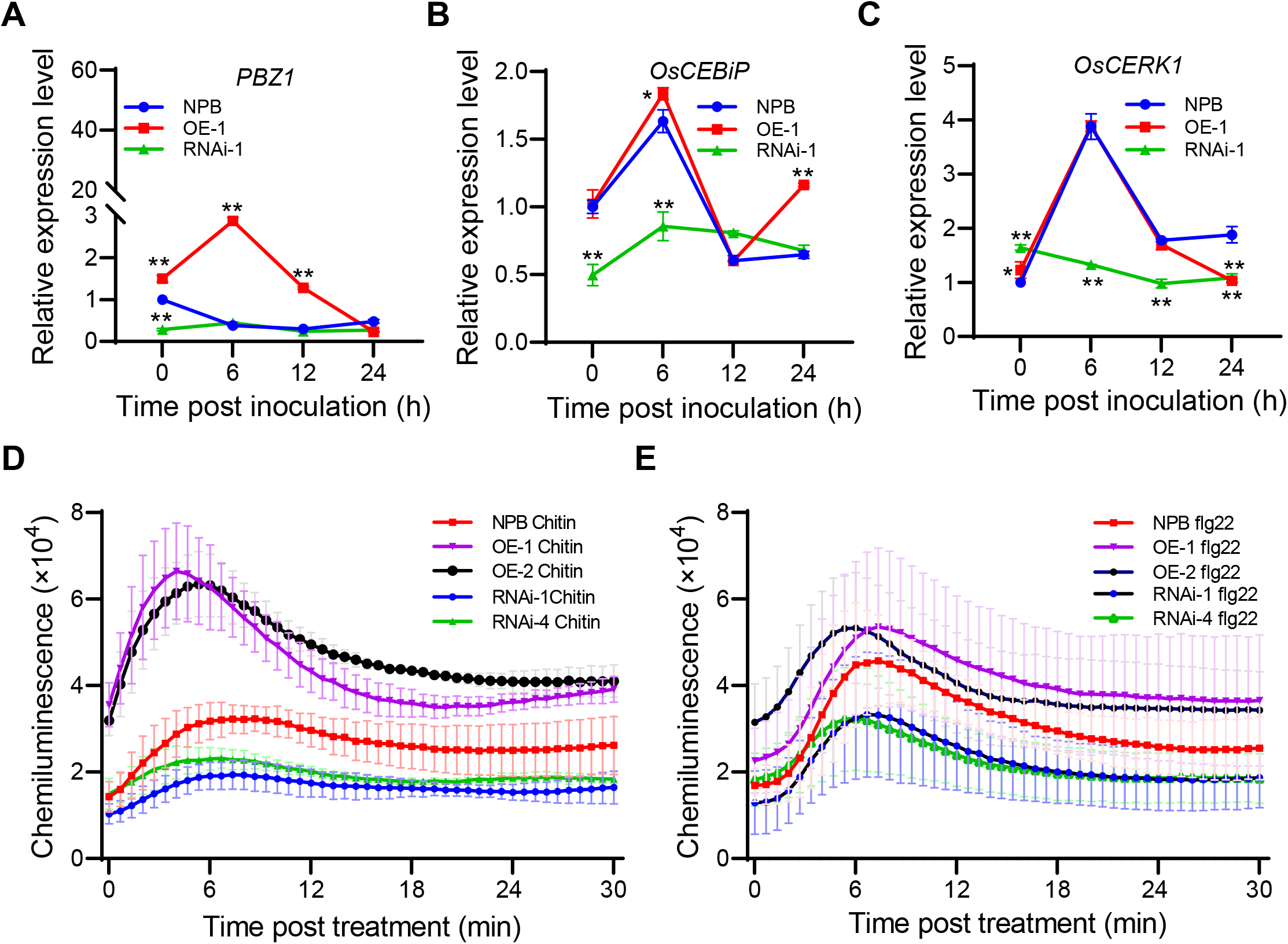
AVIN8 activates PTI signaling in rice. A-C. Expression level of defense-related genes *PBZ1* (A), *OsCEBiP* (B), *OsCERK1* (C) in *AVIN8* transgenic rice infected with *M. oryza*e strain 318-2. NPB is the wild type, OE-1 is *AVIN8* overexpression line, RNAi-1 is *AVIN8* RNA silencing line. Student *t*-test method was used for significance analysis (\**P* < 0.05; \*\**P* < 0.01). D-E. ROS accumulation dynamics of *AVIN8* transgenic rice lines after treated with PAMPs chitin (D) and flg22 (E).

We further examine AVIN8’s role in PTI signaling by measuring ROS production elicited by the PAMPs chitin and flg22. Following treatment with either PAMPs, ROS accumulation was more rapid and pronounced in *AVIN8-*OE lines compared to NPB, whereas *AVIN8-*RNAi line exhibited attenuated ROS generation (Fig. 5D, E). These results indicate that AVIN8 positively regulates PTI signaling in rice.

### AVIN8 involves in calcium-mediated immune signaling

Recent studies have shown that plant ANK proteins such as ACD6 and Lr14a function as calcium-permeable channels to mediate Ca^2+^ influx during immune activation (Chen et al., 2023; Yue et al., 2025). Because AVIN8 shares conserved residues in its ANK domain with these known channels (Fig. 2B), we investigated whether AVIN8 is similarly involved in Ca^2+^ influx regulation. Ca^2+^-elicited ROS accumulation dynamics were measured in *AVIN8* transgenic lines following treatment with exogenous Ca^2+^ and the calcium-channel inhibitor LaCl_3_. The results showed that Ca^2+^-triggered ROS production was enhanced in *AVIN8-*OE lines but attenuated in *AVIN8*-RNAi lines (Fig. 6A, B). This Ca^2+^-induced ROS burst was effectively suppressed by LaCl_3_. Furthermore, the elevated chitin-triggered ROS generation observed in *AVIN8-*OE lines was also significantly reduced upon LaCl_3_ treatment (Fig. 6C, D). These results suggest that AVIN8 likely modulates Ca^2+^ influx to regulate PTI-associated immune responses in rice.

**Figure 6.**
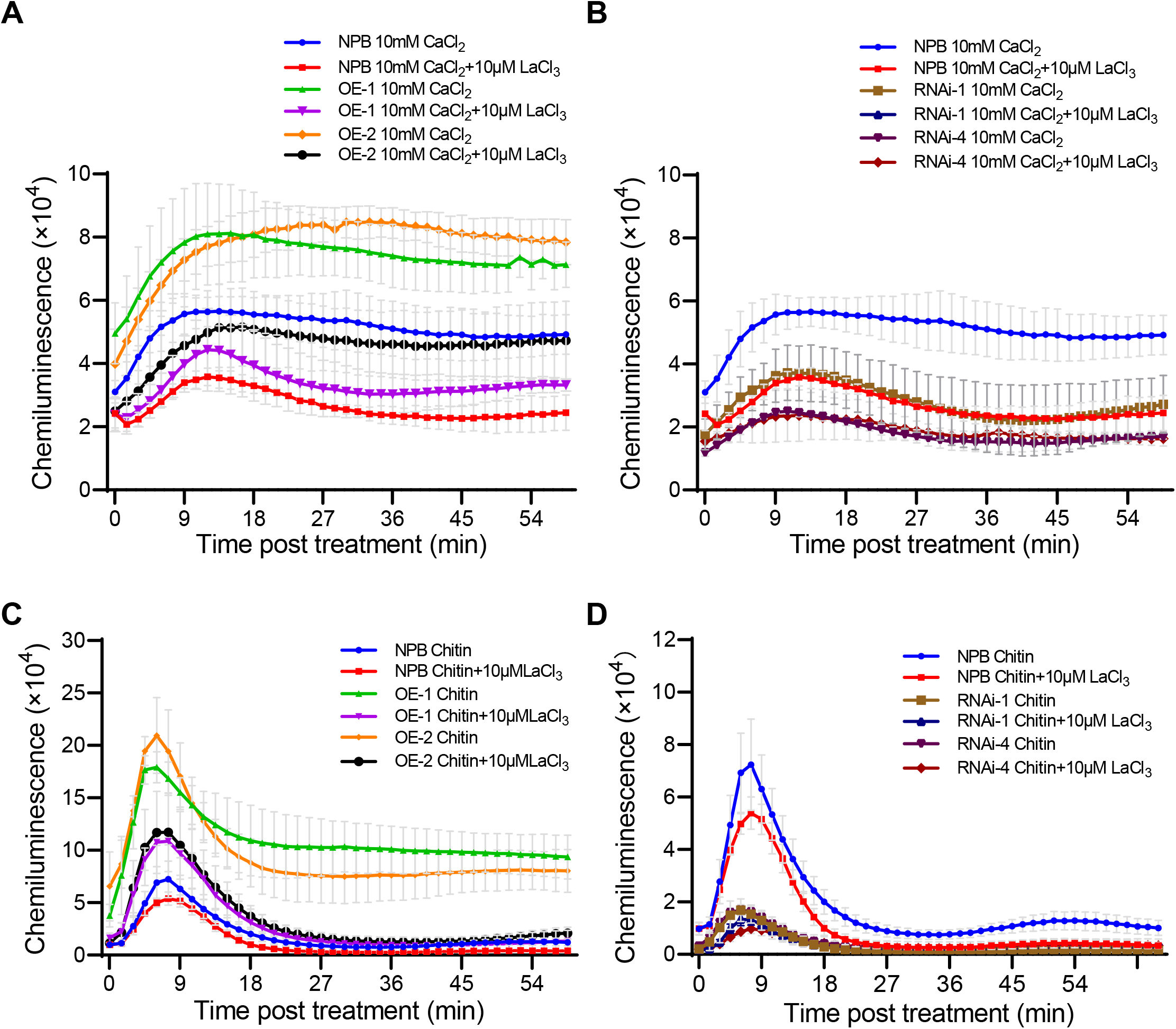
AVIN8 positively regulates calcium-elicited ROS production. A-B. ROS accumulation dynamics with the treatment of Ca^2+^ and calcium channel inhibitor LaCl_3_ in *AVIN8*-OE (A) and RNAi (B) lines. OE-1 and OE-2 are two independent *AVIN8* OE lines, RNAi-1 and RNAi-4 are two independent *AVIN8* RNAi lines. C-D. ROS accumulation dynamics with the treatments of Chitin and calcium channel inhibitor LaCl_3_ in *AVIN8*-OE (C) and RNAi (D) lines.

### AvrPii disrupts calcium-dependent immune activation

To determine whether AvrPii impairs calcium-dependent immune signaling, we evaluated Ca^2+^ and chitin-elicited ROS production in *AvrPii*-expressing rice lines following treatment with exogenous Ca^2+^ and the calcium-channel inhibitor LaCl_3_. The results revealed that both Ca^2+^ and chitin-elicited ROS bursts were significantly attenuated in *AvrPii*-expressing lines compared with the wild-type NPB (Fig. 7A, B). Moreover, LaCl_3_ treatment further suppressed ROS generation in *AvrPii* transgenic lines (Fig. 7A, B), suggesting that AvrPii interferes with Ca^2+^-dependent ROS production during immune activation.

**Figure 7.**
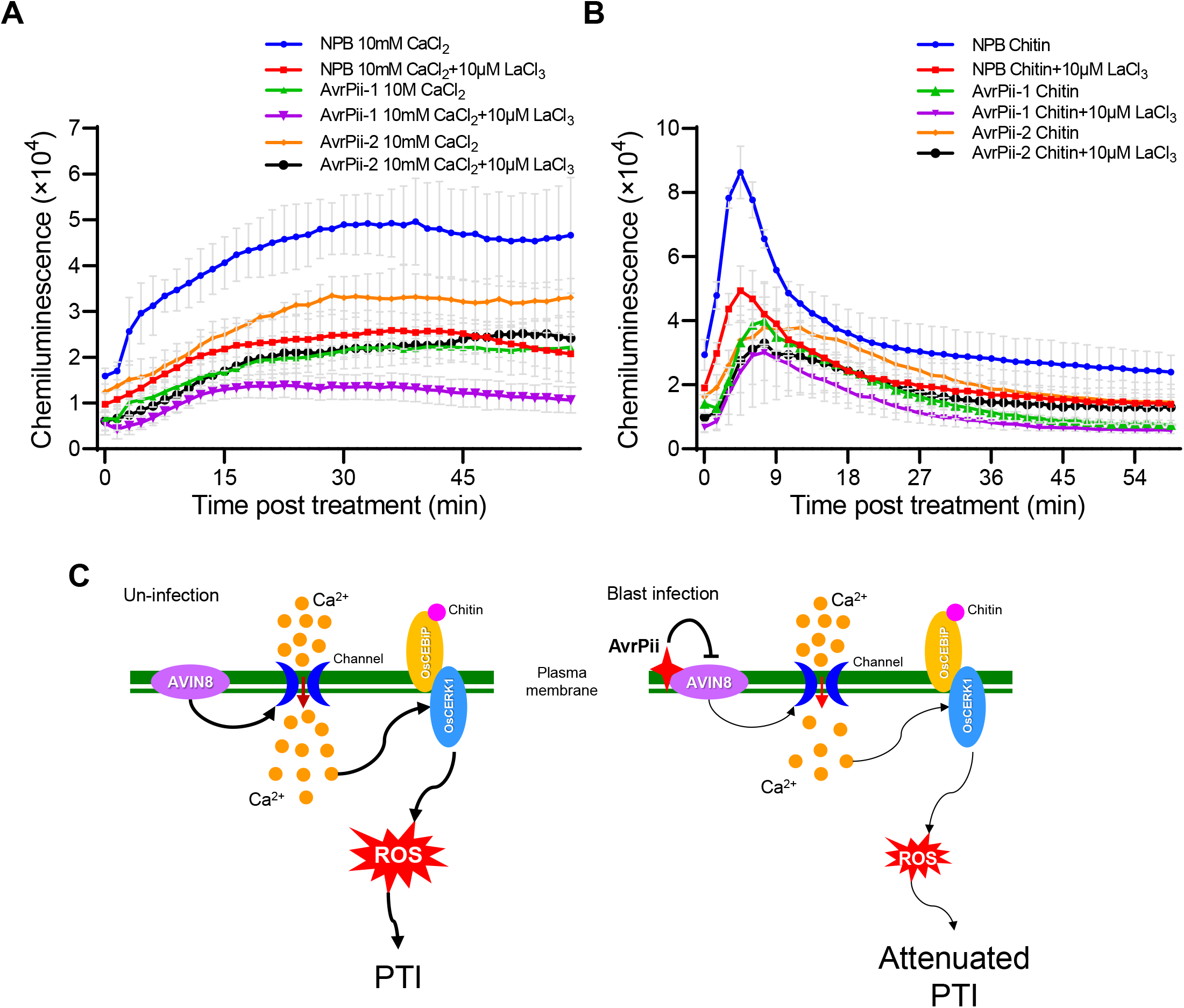
AvrPii dampens Ca^2+^-triggering ROS burst in rice. A. ROS accumulation dynamics with the treatment of Ca^2+^ and calcium channel inhibitor LaCl_3_ in *AvrPii* transgenic rice lines. AvrPii-1 and AvrPii-2 are two independent *AvrPii* ectopic expression rice lines. B. ROS accumulation dynamics with the treatment of Chitin and calcium channel inhibitor LaCl_3_ in *AvrPii* transgenic rice lines. C. Proposed work model of AvrPii-AVIN8 association-mediated immune response in rice. In the model, the membrane localized ANK protein AVIN8 function as a calcium-associated protein to promote calcium influx, thereby activating PTI immune response. When the rice blast fungal harboring the effector AvrPii infection, the secreted AvrPii physically binds with AVIN8, and suppresses AVIN8-mediated calcium influx, resulting in an inhibition of PTI signaling and promotion of blast infection.

## Discussion

In this study, we identify AVIN8 as a previously uncharacterized host target of the *M. oryzae* effector AvrPii. AVIN8 is an ANK protein that exhibits dual localization to the plasma membrane and the nuclear envelope. While plant ANK proteins have been reported to localize in diverse subcellular compartments, including the plasma membrane, nucleus, cytosol, chloroplast, mitochondria, and endoplasmic reticulum (Vo et al., 2015), dual localization to the plasma membrane and nuclear envelope has not been described. This unique localization suggests that AVIN8 may function at the interface of membrane-initiated signaling and nuclear regulatory events-a previously unrecognized feature among plant ANK proteins.

In rice, several immunity-associated ANK genes are transcriptionally up-regulated upon infection by pathogens such as *M. oryzae* and *Xanthomonas oryzae pv. Oryzae* (*Xoo*) (Vo et al., 2015). Consistently, we found that *AVIN8* expression is strongly up-regulated during *M. oryzae* infection. Notably, this induction occurs from the early stage of blast infection (0-12hpi), implicating that AVIN8 is likely a participant in the initial immune activation. Functional analyses further demonstrated that AVIN8 positively regulates blast resistance and is associated with enhanced expression of *PR* genes during early infection. Thus, these results indicate that AVIN8 acts as a positive regulator of the early immune response against *M. oryzae*.

PTI, initiated by the cell-surface receptors such as FLS2 and CEBiP/CERK1, constitutes an early defense line against pathogens (Ngou et al., 2022). In rice, the co-receptors OsCEBiP and OsCERK1 perceive fungal chitin and activate PTI signaling. Given the plasma-membrane localization, AVIN8 was hypothesized to participate in PTI. Supporting this, we observed that expression of *OsCEBiP* and *CERK1* was significantly suppressed in *AVIN8*-RNAi lines, while only *OsCEBiP* expression was up-regulated in *AVIN8*-OE lines during *M. oryzae* infection. The selective upregulation of *OsCEBiP* but not *OsCERK1* in OE lines may reflect distinct regulatory requirements for these co-receptors, or suggest that AVIN8 primarily influences the chitin perception module at the receptor level. Moreover, chitin-elicited ROS production was markedly enhanced in *AVIN8*-OE lines but attenuated in *AVIN8*-RNAi lines. These results demonstrate that AVIN8 functions as a positive regulator of chitin-triggered PTI in rice. However, how AVIN8 associates with chitin receptors OsCEBiP and OsCERK1 to trigger PTI requires further investigation.

Recent studies have revealed that plant ANK proteins such as ACD6 and Lr14a can act as calcium-permeable channels to mediate Ca^2+^ influx and promote immune activation, analogous to the human ANK channel TRPA1 (Paulsen et al., 2015; Chen et al., 2023; Yue et al., 2025). Here, we show that the ANK domain of AVIN8 shares sequence similarity with that of TRPA1, raising the possibility that AVIN8 may also participate in calcium signaling during rice immunity. Consistent with this hypothesis, our data show AVIN8 positively regulates Ca^2+^-elicited ROS production, in which overexpressing *AVIN8* in rice significantly enhanced exogenous Ca^2+^-elicited ROS burst, whereas silencing *AVIN8* impaired Ca^2+^-induced ROS accumulation. Furthermore, both Ca^2+^- and chitin-elicited ROS generation were suppressed by the calcium channel inhibitor LaCl_3_. These results indicate that AVIN8 functions as a positive modulator of calcium-mediated immune activation. Whether AVIN8 itself form a functional Ca^2+^ channel remains to be determined. Electrophysiological analyses using heterologous systems (e.g., *Xenopus oocytes*, HEK293T cells) will be essential to determine whether AVIN8 exhibits intrinsic channel activity.

Calcium influx and ROS burst are intimately interconnected signaling events that driven early immune activation (Köster et al., 2022). Our data demonstrate that the membrane-localized ANK protein AVIN8 positively regulates Ca^2+^ influx and ROS production. Treatment with the calcium-channel inhibitor LaCl_3_ markedly reduced both Ca^2+^- and chitin-triggered ROS bursts, indicating that AVIN8 is required for Ca^2+^ dependent ROS generation during PTI in rice. Given that AVIN8 is directly targeted by the blast fungal effector AvrPii, we hypothesized that AvrPii may interfere with AVIN8-mediated Ca^2+^ signaling. Indeed, Ca^2+^- and chitin-elicited ROS production was significantly attenuated in *AvrPii-* expressing rice lines, and LaCl_3_ treatment further enhanced this suppression. Therefore, these results demonstrate that *M. oryzae* AvrPii disrupts PTI signaling, at least in part, by impairing AVIN8-mediated Ca^2+^ influx and ROS generation.

Although *M. oryzae* effectors are known to target diverse signaling components in both apoplastic and cytoplasmic compartments to manipulate host immunity (Oliveira-Garcia et al., 2024), direct targeting of host membrane proteins as an immune suppression strategy remains rarely documented. To date, only the effector AvrPiz-t has been shown to interact with the plasma membrane K^+^ channel protein OsAKT1 disrupting ion homeostasis and immunity (Shi et al., 2018). Unlike AvrPiz-t, which targets a K^+^ channel, we demonstrate that AvrPii directly targets a Ca^2+^ signaling-associated ANK membrane protein AVIN8 to interfere with Ca^2+^ signaling and PTI responses, providing new evidence that direct targeting of host membrane components represents an emerging strategy by filamentous pathogens to subvert plant immunity. Collectively, our work uncovers a distinct virulence mechanism in which *M. oryzae* effector AvrPii functions as a suppressor of calcium-influx by exploiting the membrane-localized calcium regulator AVIN8 to disarm PTI in rice (Fig. 7C).

## Acknowledgments

We thank Mr. Wei Li and Mr. Xingguo Liu for their assistance in laboratory work and field managements, and thank Dr. Feng He and Dr. Guo-Liang Wang for their assistance on the yeast two-hybrid screening. This work was supported by the grants from the National Natural Science Foundation of China (#31401726 and #31972256), Natural Science Foundation of Hunan Province (#2025JJ50127), Scientific Research Project of the Hunan Provincial Education Department in China (#24A0165) and Graduate Research Innovation Project in Huan Province of China (#CX20240637).

## Author contributions

JLL conceived the project and supervised the research. WY performed the experiments, analyzed the data, and drafted the manuscript. FHY, XLZ, QRC, JLL, YYZ, CYX, BH, DDZ, SSJ performed partial experiments and data collection. XLL co-supervised the investigation, discussed the data and manuscript revision. JLL revised and finalized the manuscript.

## Supplemental Figure Legends

**Figure S1. Sequence similarity of the ANK domains of AVIN8 and known calcium-channel proteins ACD6, Lr14a and TRPA1**

A. Sequence similarity of the ANK domains between AVIN8 and known calcium-channel proteins ACD6, Lr14a and TRPA1. B. Phylogenetic relationship of the ANK domains between AVIN8 and known calcium-channel proteins ACD6, Lr14a and TRPA1.

**Figure S2. Construction of *AVIN8* transgenic rice**

A. AVIN8 expression level in *AVIN8*-OE transgenic rice lines. B. *AVIN8* dsRNA position and sequence in the 3’ UTR region for RNAi constructs making. C. *AVIN8* expression level in AVIN8-RNAi transgenic rice lines.

## References

Cao, H., Glazebrook, J., Clarke, J.D., Volko, S., Dong, X. (1997) The Arabidopsis NPR1 gene that controls systemic acquired resistance encodes a novel protein containing ankyrin repeats. Cell, 88(1):57–63.

Chen, J., Li, L., Kim, J.H., Neuhäuser, B., Wang, M., Thelen, M., Hilleary, R., Chi, Y., Wei, L., Venkataramani, K., Exposito-Alonso, M., Liu, C., Keck, J., Barragan, A.C., Schwab, R., Lutz, U., Pei, Z.M., He, S.Y., Ludewig, U., Weigel, D., Zhu, W. (2023) Small proteins modulate ion-channel-like ACD6 to regulate immunity in Arabidopsis thaliana. Mol Cell, 83(23):4386-4397.e9.

Chen, K., Zhuang, Y., Chen, H., Lei, T., Li, M., Wang, S., Wang, L., Fu, H., Lu, W., Bohra, A., Lai, Q., Xu, X., Garg, V., Barmukh, R., Ji, B., Zhang, C., Pandey, M.K., Tang, R., Varshney, R.K., Zhuang, W. A. (2025) Ralstonia effector RipAU impairs peanut AhSBT1.7 immunity for pathogenicity via AhPME-mediated cell wall degradation. Plant J, 121(2): e17210.

Chen, S., Songkumarn, P., Liu, J., Wang, G.L. (2009) A versatile zero background T-vector system for gene cloning and functional genomics. Plant Physiol, 150(3):1111–21.

Chern, M., Fitzgerald, H.A., Canlas, P.E., Navarre, D.A., Ronald, P.C. (2005) Overexpression of a rice NPR1 homolog leads to constitutive activation of defense response and hypersensitivity to light. Mol Plant Microbe Interact, 18(6):511–20.

De la Concepcion, J.C., Fujisaki, K., Bentham, A.R., Mireles, N.C., de Medina Hernandez, V.S., Shimizu, M., Lawson, D.M., Kamoun, S., Terauchi, R., Banfield, M.J. (2022) A blast fungus zinc-finger fold effector binds to a hydrophobic pocket in host Exo70 proteins to modulate immune recognition in rice. Proc Natl Acad Sci U S A, 19 (43): e2210559119.

Fujisaki, K., Abe, Y., Ito, A., Saitoh, H., Yoshida, K., Kanzaki, H., Kanzaki, E., Utsushi, H., Yamashita, T., Kamoun, S., Terauchi, R. (2015) Rice Exo70 interacts with a fungal effector, AVR-Pii, and is required for AVR-Pii-triggered immunity. Plant J, 83(5):875–87.

Gong, B.Q., Wang, F.Z., Li, J.F. (2020) Hide-and-Seek: Chitin-triggered plant immunity and fungal counterstrategies. Trends Plant Sci, 25(8):805–816.

Goodin, M.M., Dietzgen, R.G., Schichnes, D., Ruzin, S., Jackson, A.O. (2002) pGD vectors: versatile tools for the expression of green and red fluorescent protein fusions in agroinfiltrated plant leaves. Plant J, 31(3):375–83.

Han, J., Wang, X., Wang, F., Zhao, Z., Li, G., Zhu, X., Su, J., Chen, L. (2021) The fungal effector Avr-Pita suppresses innate immunity by increasing COX activity in rice mitochondria. Rice (N Y), 14(1):12.

Hu, Y., Ding, Y., Cai, B., Qin, X., Wu, J., Yuan, M., Wan, S., Zhao, Y., Xin, X.F. (2022) Bacterial effectors manipulate plant abscisic acid signaling for creation of an aqueous apoplast. Cell Host Microbe, 30(4):518-529.e6.

Huang, X., Yao, W., Chen, Q., Lin, J., Huang, J., Zou, Y., Guo, C., He, B., Yuan, X., Xu, C., Liu, X., Xiao, Y., Wu, J., Liu, J. (2025) A century of advances in molecular genetics and breeding for sustainable resistance to rice blast disease. Theor Appl Genet, 138(7):174.

Jiang, Y., Chen, X., Ding, X., Wang, Y., Chen, Q., Song, W.Y. (2013) The XA21 binding protein XB25 is required for maintaining XA21-mediated disease resistance. Plant J, 73(5):814–23.

Jones, J.D.G., Staskawicz, B.J., Dangl, J.L. (2024) The plant immune system: From discovery to deployment. Cell, 187(9):2095–2116

Kim, S., Kim, C.Y., Park, S.Y., Kim, K.T., Jeon, J., Chung, H., Choi, G., Kwon, S., Choi, J., Jeon, J., Jeon, J.S., Khang, C.H., Kang, S., Lee, Y.H. (2020) Two nuclear effectors of the rice blast fungus modulate host immunity via transcriptional reprogramming. Nat Commun, 11(1):5845.

Kitazawa, Y., Iwabuchi, N., Maejima, K., Sasano, M., Matsumoto, O., Koinuma, H., Tokuda, R., Suzuki, M., Oshima, K., Namba, S., Yamaji, Y. (2022) A phytoplasma effector acts as a ubiquitin-like mediator between floral MADS-box proteins and proteasome shuttle proteins. Plant Cell, 34(5):1709–1723.

Köster, P., DeFalco, T.A., Zipfel C. (2022) Ca^2+^ signals in plant immunity. EMBO J., 41(12):e110741.

Li, B., Zhou, X., Yao, W., Lin, J., Ding, X., Chen, Q., Huang, H., Chen, W., Huang, X., Pan, S., Xiao, Y., Liu, J., Liu, X., Liu, J. (2024) NADP-malic enzyme OsNADP-ME2 modulates plant height involving in gibberellin signaling in rice. Rice (N Y), 17(1):52.

Li, Y., Jin, K., Ghosh, S., Devarakonda, P., Carlson, K., Davis, A., Stewart, K.A.V., Cammett, E., Rossi, P.P., Setlow, B., Lu, M., Setlow, P., Hao, B. (2014) Structural and functional analysis of the GerD spore germination protein of Bacillus species. J Mol Biol, 426(9):1995–2008.

Liu, C., Han, L.B., Wen, Y., Lu, C., Deng, B., Liu, Z., Deng, X., Shen, N., Tang, D., Li, Y.B. (2025) The Magnaporthe oryzae effector MoBys1 suppresses rice immunity by targeting OsCAD2 to manipulate host jasmonate and lignin metabolism. New Phytol, 246(1):280–297.

Liu, X., Gao, Y., Guo, Z., Wang, N., Wegner, A., Wang, J., Zou, X., Hu, J., Liu, M., Zhang, H., Zheng, X., Wang, P., Schaffrath, U., Zhang, Z. (2022) MoIug4 is a novel secreted effector promoting rice blast by counteracting host OsAHL1-regulated ethylene gene transcription. New Phytol, 235(3):1163–1178.

Liu, X., Yu, Y., Yao, W., Yin, Z., Wang, Y., Huang, Z., Zhou, J.Q., Liu, J., Lu, X., Wang, F., Zhang, G., Chen, G., Xiao, Y., Deng, H., Tang, W. (2023) CRISPR/Cas9-mediated simultaneous mutation of three salicylic acid 5-hydroxylase (OsS5H) genes confers broad-spectrum disease resistance in rice. Plant Biotechnol J, 21(9):1873–1886.

Lu, H., Rate, D.N., Song, J.T., Greenberg, J.T. (2003) ACD6, a novel ankyrin protein, is a regulator and an effector of salicylic acid signaling in the Arabidopsis defense response. Plant Cell, 5(10):2408–20.

Martinez-D’Alto, A., Yan, X., Detomasi, T.C., Sayler, R.I, Thomas, W.C., Talbot, N.J., Marletta. M.A. (2023) Characterization of a unique polysaccharide monooxygenase from the plant pathogen Magnaporthe oryzae. Proc Natl Acad Sci U S A, 120(8): e2215426120.

Mentlak, T.A., Kombrink, A., Shinya, T., Ryder, L.S., Otomo, I., Saitoh, H., Terauchi, R., Nishizawa, Y., Shibuya, N., Thomma, B.P.H.J., Talbot, N.J. (2012) Effector-mediated suppression of chitin-triggered immunity by Magnaporthe oryzae is necessary for rice blast disease. Plant Cell, 24(1):322–35

Mou, S., Liu, Z., Guan, D., Qiu, A., Lai, Y., He, S. (2013) Functional analysis and expressional characterization of rice ankyrin repeat-containing protein, OsPIANK1, in basal defense against Magnaporthe oryzae attack. PLoS One, 8(3):e59699.

Ngou, B.P.M., Ding, P., Jones, J.D.G. (2022) Thirty years of resistance: Zig-zag through the plant immune system. Plant Cell, 34(5):1447–1478.

Ning, N., Xie, X., Yu, H., Mei, J., Li, Q., Zuo, S., Wu, H., Liu, W., Li. Z. (2022) Plant peroxisome-targeting effector MoPtep1 is required for the virulence of Magnaporthe oryzae. Int J Mol Sci, 23(5):2515.

Oikawa, K., Fujisaki, K., Shimizu, M., Takeda, T., Nemoto, K., Saitoh, H., Hirabuchi, A., Hiraka, Y., Miyaji, N., Białas, A., Langner, T., Kellner, R., Bozkurt, T.O., Cesari, S., Kroj, T., Banfield, M.J., Kamoun, S., Terauchi, R. (2024) The blast pathogen effector AVR-Pik binds and stabilizes rice heavy metal-associated (HMA) proteins to co-opt their function in immunity. PLoS Pathog, 20(11): e1012647.

Oliveira-Garcia, E., Yan, X., Oses-Ruiz, M., de Paula, S., Talbot, N.J. (2024) Effector-triggered susceptibility by the rice blast fungus Magnaporthe oryzae. New Phytol, 241(3):1007–1020.

Park, C.H., Chen, S., Shirsekar, G., Zhou, B., Khang, C.H., Songkumarn, P., Afzal, A.J., Ning, Y., Wang, R., Bellizzi, M., Valent, B., Wang. G.L. (2012) The Magnaporthe oryzae effector AvrPiz-t targets the RING E3 ubiquitin ligase APIP6 to suppress pathogen-associated molecular pattern-triggered immunity in rice. Plant Cell, 24(11):4748–62.

Paulsen, C.E., Armache, J.P., Gao, Y., Cheng, Y., Julius, D. (2015) Structure of the TRPA1 ion channel suggests regulatory mechanisms. Nature, 520(7548):511–7.

Peck, S.C., Nühse, T.S., Hess, D., Iglesias, A., Meins, F., Boller T. (2001) Directed proteomics identifies a plant-specific protein rapidly phosphorylated in response to bacterial and fungal elicitors. Plant Cell 13:1467–1475.

Schmid, A.B, Lagleder, S., Gräwert, M.A., Röhl, A., Hagn, F., Wandinger, S.K., Cox, M.B., Demmer, O., Richter, K., Groll, M., Kessler, H., Buchner, J. (2012) The architecture of functional modules in the Hsp90 co-chaperone Sti1/Hop. EMBO J, 31(6):1506–17.

Shi, X., Long, Y., He, F., Zhang, C., Wang, R., Zhang, T., Wu, W., Hao, Z., Wang, Y., Wang, G.L., Ning, Y. (2018) The fungal pathogen Magnaporthe oryzae suppresses innate immunity by modulating a host potassium channel. PLoS Pathog, 14(1): e1006878.

Singh, R., Dangol, S., Chen, Y., Choi, J., Cho, Y.S., Lee, J.E., Choi, M.O., Jwa, N.S. (2016) Magnaporthe oryzae effector AVR-Pii helps to establish compatibility by inhibition of the rice NADP-Malic enzyme resulting in disruption of oxidative burst and host innate immunity. Mol Cells, 39(5):426–38.

Sun, Q., Fan, M., Qiu, H., Zuo, Y., Lin, T., Xu, M., Nie, J., Wu, J., Zhou, J., Yu, R., Liu, L., Tian, Z. (2025) Phytophthora RxLR effector Pi18609 suppresses host immunity by promoting turnover of a positive immune regulator StBBX27. Plant J, 121(5): e70074.

Vo, K.T.X., Lee, S.K., Halane, M.K., Song, M.Y., Hoang, T.V., Kim, C.Y., Park, S.Y., Jeon, J., Kim, S.T., Sohn, K.H., Jeon, J.S. (2019) Pi5 and Pii Paired NLRs are functionally exchangeable and confer similar disease resistance specificity. Mol Cells, 42(9):637–645.

Waadt, R., Schmidt, L.K., Lohse, M., Hashimoto, K., Bock, R., Kudla, J. (2008) Multicolor bimolecular fluorescence complementation reveals simultaneous formation of alternative CBL/CIPK complexes in planta. Plant J, 56(3):505–16.

Wang, W., Cheng, H.Y., Zhou, J.M. (2024) New insight into Ca^2+^-permeable channel in plant immunity. J Integr Plant Biol, 66(3):623–631.

Wang, Y., Pruitt, R.N., Nürnberger, T., Wang, Y. (2022) Evasion of plant immunity by microbial pathogens. Nat Rev Microbiol, 20(8):449–464.

Wang, Y.S., Pi, L.Y., Chen, X., Chakrabarty, P.K., Jiang, J., Leon, A.L.D., Liu, G.Z., Li, L., Benny, U., Oard, J., Ronald, P.C., Song, W.Y. (2006) Rice XA21 binding protein 3 is a ubiquitin ligase required for full Xa21-mediated disease resistance. Plant Cell 18(12):3635-46.

Wang, Z., Zhong, G., Zhang, B., Xie, Y., Gan, Y., Tang, D., Wang, W. (2026) Rice blast pathogen effector AvrPib compromises disease resistance by targeting Raf-like protein kinase OsMAPKKK72 to inhibit MAPK signaling. J Integr Plant Biol, 68(2):486–501.

Waszczak, C., Carmody, M., and Kangasjärvi, J. (2018) Reactive Oxygen Species in Plant Signaling. Annu. Rev. Plant Biol, 69:5.1-5.28

Wu, M., Zhao, Y., Yang, J., Yang, F., Dai, Y., Wang, Q., Chen, C., Chu. X. (2025) The role of ankyrin repeat-containing proteins in epigenetic and transcriptional regulation. Cell Death Discov, 11(1):232.

Xu, G., Moeder, W., Yoshioka, K., Shan, L. (2022) A tale of many families: calcium channels in plant immunity. Plant Cell, 34(5):1551–1567.

Yang, Y., Zhang, Y., Ding, P., Johnson, K., Li, X., Zhang, Y. (2012) The ankyrin-repeat transmembrane protein BDA1 functions downstream of the receptor-like protein SNC2 to regulate plant immunity. Plant Physiol, 159(4):1857–65.

Yue, L., Wang, L., Neuhäuser, B., Zhang, S., Herren, G., Jung, E., Kim, G., Goto, Y., Fernández-Fernández, Á.D., Heuberger, M., Ludewig, U., Zipfel, C., Keller, B. (2025) Cytoplasmic calcium influx mediated by Lr14a regulates stomatal immunity against leaf rust in wheat. Curr Biol, 35(23):5750-5761.e4.

Zhang, F., Fang, H., Wang, M., He, F., Tao, H., Wang, R., Long, J., Wang, J., Wang, G.L., Ning, Y. (2022) APIP5 functions as a transcription factor and an RNA-binding protein to modulate cell death and immunity in rice. Nucleic Acids Res, 50(9):5064–5079.

Zhang, S., Wang, L., Jiang, H., Sun, G., Xia, Y., Wu, J., Chen, X., Wang, L., Liu, T., Ouyang, H., Chen, X., Wang, Y., Wang, Y. (2025) A conserved Phytophthora apoplastic trypsin-like serine protease targets the receptor-like kinase BAK1 to dampen plant immunity. Nat Plants, 11(7):1401–1415.

Zhang, X., Li, D., Zhang, H., Wang, X., Zheng, Z., Song, F. (2010) Molecular characterization of rice OsBIANK1, encoding a plasma membrane-anchored ankyrin repeat protein, and its inducible expression in defense responses. Mol Biol Rep, 37:653–660.

Zheng, P., Yuan, P., Fang, N., Ma, X., Jiang, L., Zeng, Q., Han, D., Kang, Z., Liu, J. (2025) Stripe rust fungi hijack a host transcriptional repressor to induce TaSWEET14d for enhanced sugar uptake in wheat. Sci Adv,11(28): eadv1760.

