## Supplemental figures and tables for "The *Magnaporthe oryzae* Effector AvrPii Attenuates Rice Immunity by Targeting the Calcium-Associated ANK Protein AVIN8"

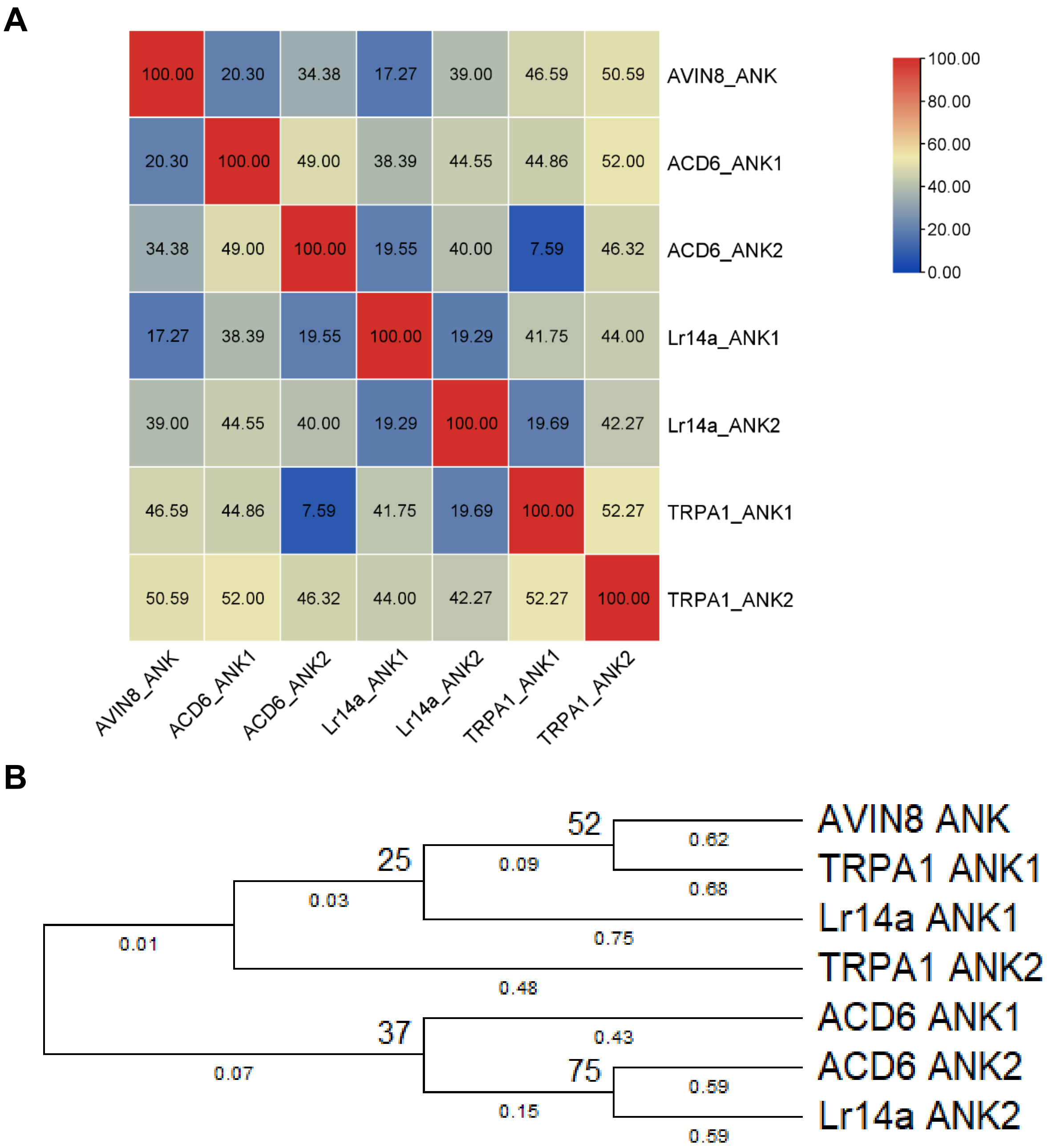


**Figure S1. Sequence similarity of the ANK domains of AVIN8 and known calcium-channel proteins ACD6, Lr14a and TRPA1**

A. Sequence similarity of the ANK domains between AVIN8 and known calcium-channel proteins ACD6, Lr14a and TRPA1. B. Phylogenetic relationship of the ANK domains between AVIN8 and known calcium-channel proteins ACD6, Lr14a and TRPA1.

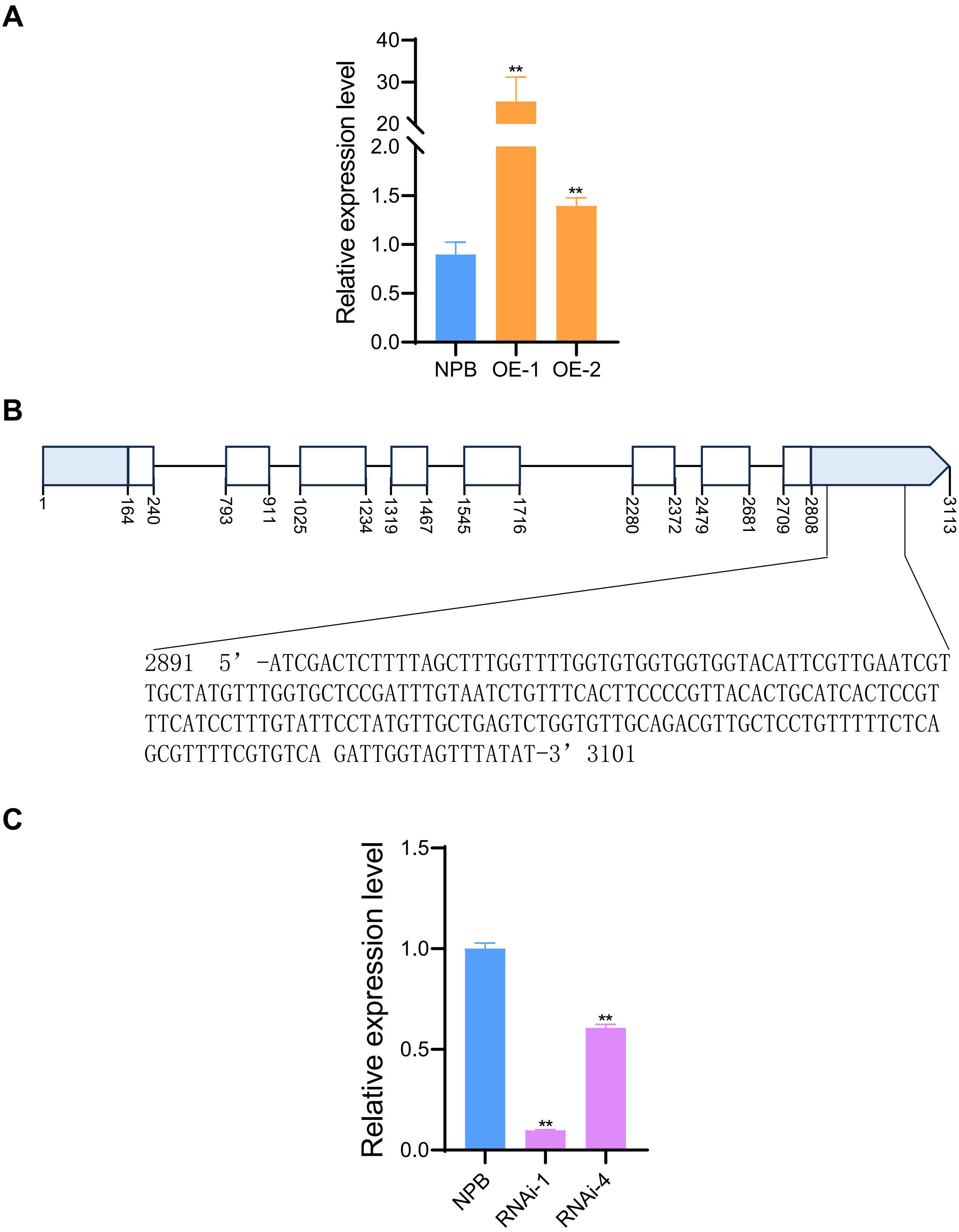


**Figure S2. Construction of *AVIN8* transgenic rice**

A. *AVIN8* expression level in *AVIN8*-OE transgenic rice lines. B. *AVIN8* dsRNA position and sequence in the 3’ UTR region for RNAi constructs making. C. *AVIN8* expression level in *AVIN8*-RNAi transgenic rice lines.

### Supplemental Tables

Table S1 The primer pairs used in this study

| Primer name | Primer sequence | Goals |
|---|---|---|
| AVIN8FL-F | 5'-TATGGAAGGTATGATGATGATTTGTTC-3' | *AVIN8* full length cDNA cloning |
| AVIN8FL-R | 5'-TTATAGGAAAGCGTCCATCTCG-3' | |
| AVIN8Y2HF-SmaI | ATAA C CCG GGT ATGGAAGACCAGAAGAAAAATGC | *AVIN8* Y2H AD vector making |
| AVIN8Y2HR-NotI | CATA GC GGC CGC TTATAGGAAAGCGTCCATCTCG | |
| AvrPiiF-Y2HSalI | TCGAGG TCG ACC ATGCAACTTTCCAAAATTAC | *AvrPii* Y2H BD vector making |
| AvrPiiR-Y2HNotI | CTTAGC GGC CGC TTAGTTGCATTTATGATTAA | |
| AvrPiiBiFCFLF-BamHI | ATAA GGA TCC ATGCAACTTTCCAAAATTAC | *AvrPii* C-YFP vector makaing |
| AvrPiiBiFCFLR-XhoI | TCAT CTC GAG TTAGTTGCATTTATGATTAA | |
| AVIN8BiFCFLF-SalI | ATAA GTC GAC ATGGAAGGTATGATGATGATTTGTTC | *AVIN8* N-YFP vector making |
| AVIN8BiFCFLR-KpnI | TCAT GGT ACC TTATAGGAAAGCGTCCATCTCG | |
| AVIN8GDGFLF-BglII | TCGC AGA TCT ATGGAAGGTATGATGATGATTTGTTC | *AVIN8* GFP fusion vector making |
| AVIN8GDGFLR-SalI | ACGC GTC GAC TTATAGGAAAGCGTCCATCTCG | |
| AVIN8Ri-QF | GGCTCGTCGCTTTACAGTTC | *AVIN8* qPCR detection in RNAi lines |
| AVIN8Ri-QR | ATCGGAGCACCAAACATAGC | |
| AVIN8OX-QF | CGTGTGGTTACGGTGAGTTG | *AVIN8* qPCR detection in OX lines |
| AVIN8OX-QR | TATCCAGCAGCGTAATGCAG | |
| Ubq-QF | CGCAAGAAGAAGTGTCCTCA | *Ubiquitin* gene qPCR detection |
| Ubq-QR | GGGAGATAACAACGGAAGCA | |
| PBZ1-QF | CCCTGCCGAATACGCCTAA | *PBZ1* gene qPCR detection |
| PBZ1-QR | CTCAAACGCCACGAGAATTTG | |
| OsCEBiP-QF | ATGGAACGCTGAAGCTTGGTGAGA | *OsCEBiP* gene qPCR detection |
| OsCEBiP-QR | CTCATCCTCTAAAGAACAGAGTCA | |
| OsCERK1-QF | CCTATTGATTCCATTCTCAAGC | *OsCERK1* gene qPCR detection |
| OsCERK1-QR | GGTTCTCATACAGGTTGTTCA | |
